# Abelson-induced phosphorylation of TACC3 modulates its interaction with microtubules and affects its impact on axon outgrowth and guidance

**DOI:** 10.1101/2020.02.14.948448

**Authors:** Burcu Erdogan, Riley M. St. Clair, Garrett M. Cammarata, Timothy Zaccaro, Bryan A. Ballif, Laura Anne Lowery

## Abstract

Axon guidance is a critical process in forming the connections between a neuron and its target. The growth cone steers the growing axon towards the appropriate direction by integrating extracellular guidance cues and initiating intracellular signal transduction pathways downstream of these cues. The growth cone generates these responses by remodeling its cytoskeletal components. Regulation of microtubule dynamics within the growth cone is important for making guidance decisions. TACC3, as a microtubule plus-end binding protein, modulates microtubule dynamics during axon outgrowth and guidance. We have previously shown that embryos depleted of TACC3 displayed spinal cord axon guidance defects, while TACC3-overexpressing spinal neurons showed increased resistance to Slit2-induced growth cone collapse. Here, in order to investigate the mechanism behind TACC3-mediated axon guidance, we studied the importance of tyrosine phosphorylation induced by Abelson tyrosine kinase. We find that the phosphorylatable tyrosines within the TACC domain are important for the microtubule plus-end tracking behavior of TACC3. Moreover, TACC domain phosphorylation impacts axon outgrowth and guidance, and it also regulates microtubule extension into the growth cone periphery. Together, our results suggest that phosphorylation of TACC3 is a key regulatory mechanism by which TACC3 controls axon outgrowth and pathfinding decisions of neurons during embryonic development.

## Introduction

Regulation of cytoskeletal dynamics within the growth cone is essential for growth cone motility and navigation as the axon travels to its target (reviewed in Lowery and Van Vactor 2008). Guidance molecules that the growth cone encounters during its trip can act as repellent or attractant depending on the time, location and the signal composition of the environment that the growth cone passes. Integration and interpretation of these signals relies on signaling cascades that are initiated downstream of guidance cue receptors which will ultimately converge upon cytoskeletal elements for their rearrangements and control of growth cone motility.

Guidance signals are not homogeneously presented to the growth cone *in vivo*. While the growth cone might be exposed to repellent signals on one side, it can be exposed to attractant signals on the other side, which necessitates the asymmetric reorganization of the underlying cytoskeleton. In order to manage this asymmetric regulation, signals received by guidance cue receptors must be processed locally and immediately downstream of the site where the signal is received without necessarily leading to a global response. For example, repulsive guidance cues can cause a global growth cone collapse when they are bath-applied, whereas their local application causes collapse on the side that the protein is received, which results in growth cone steering away from the source of the signal (Buck and Zheng, 2002).

The interaction between guidance cue receptors and downstream targets is required for the growth cone’s directional movement. Microtubule plus-end tracking proteins (+TIPs), due to their localization close to the growth cone periphery, are potential targets for guidance signals. Their interaction with microtubules at the plus-ends is important for regulating microtubule growth dynamics and coordination of signal exchange between the growth cone periphery and the central domain, which is critical to the growth cone’s directional movement.

The interaction between +TIPs and microtubules can be modulated by guidance signals and their downstream intracellular signaling events. Phosphorylation of +TIPs is one such event that has been shown to modulate +TIP affinity for microtubules. For example, the affinity of CLASP for microtubules has been shown to be regulated differentially in the growth cone depending on its phosphorylation status by GSK3 kinase. Increased microtubule lattice binding activity of CLASP, as a result of GSK3 inhibition, results in axon growth inhibition through inhibition of microtubule advance into the growth cone periphery. On the other hand, plus-end binding of CLASP, as a result of GSK3 activity, promotes axon outgrowth via stabilization of microtubules (Hur et al., 2011). In addition to GSK3, CLASP has also been identified as a direct target of Abelson (Abl) tyrosine kinase (Engel et al., 2014; Lee et al., 2004). Further examination of the interaction between CLASP and Abl in *Xenopus* spinal neurons identified CLASP as a target for Abl phosphorylation and showed that phosphorylation can affect CLASP localization in neuronal growth cones (Engel et al., 2014).

Similar to CLASP, several +TIPs have been implicated to be involved in regulation of microtubule dynamics during directional movement of cells and growth cones (Bearce et al., 2015). However, only a few of them have been studied and implicated as a target for guidance cue-initiated intracellular signals (Bearce et al., 2015). We have previously characterized a microtubule plus-end tracking function for TACC3 in neuronal growth cones and showed that TACC3 overexpression enhances microtubule growth dynamics and promotes axon outgrowth (Nwagbara et al., 2014). Additionally, we have proposed a function for TACC3 during axon guidance. Reducing levels of TACC3 resulted in disorganized axon elongation of neural tube neurons in *Xenopus laevis* embryos and its overexpression mitigated growth cone collapse induced by bath-applied Slit2 in cultured neural tube explants (Erdogan et al., 2017). To further investigate the mechanism by which TACC3 overexpression exerts this opposing role against Slit2 activity, we became interested in looking at potential phosphorylation events that might target TACC3. Our previous studies highlighted a possible genetic interaction network between TACC3, its microtubule polymerase interactor XMAP215, and Abelson tyrosine kinase (Abl) (Lowery et al., 2010). Thus, we become interested in testing whether TACC3 could be a target for Abl phosphorylation downstream of Slit2 and whether its phosphorylation status would alter the interaction of TACC3 with microtubules as well as its impact on axon outgrowth and guidance.

## Results

### Abelson kinase induces phosphorylation of TACC3

To investigate whether Abelson tyrosine kinase (Abl) induces phosphorylation of TACC3, we co-expressed GFP-TACC3 and Abl in HEK293 cells. Tyrosine phosphorylation of TACC3, following GFP immunoprecipitation of GFP-TACC3, was observed with 4G10, an antibody that specifically labels phosphorylated tyrosine residues (Fig. 1A, lane 4). Phosphorylation of TACC3 was only observed when it was co-expressed with Abl (Fig. 1A, lane 3 vs lane 4). Additionally, this phosphorylation specifically happened with the tyrosine kinase Abl, since Fyn, which is another tyrosine kinase, did not induce TACC3 tyrosine phosphorylation (Fig. 1A, lane 6). Although we did not determine whether the induced tyrosine phosphorylation was a result of direct interaction between Abl and TACC3, our data show that Abl expression can induce phosphorylation of tyrosine residues on TACC3.

**Figure 1.**
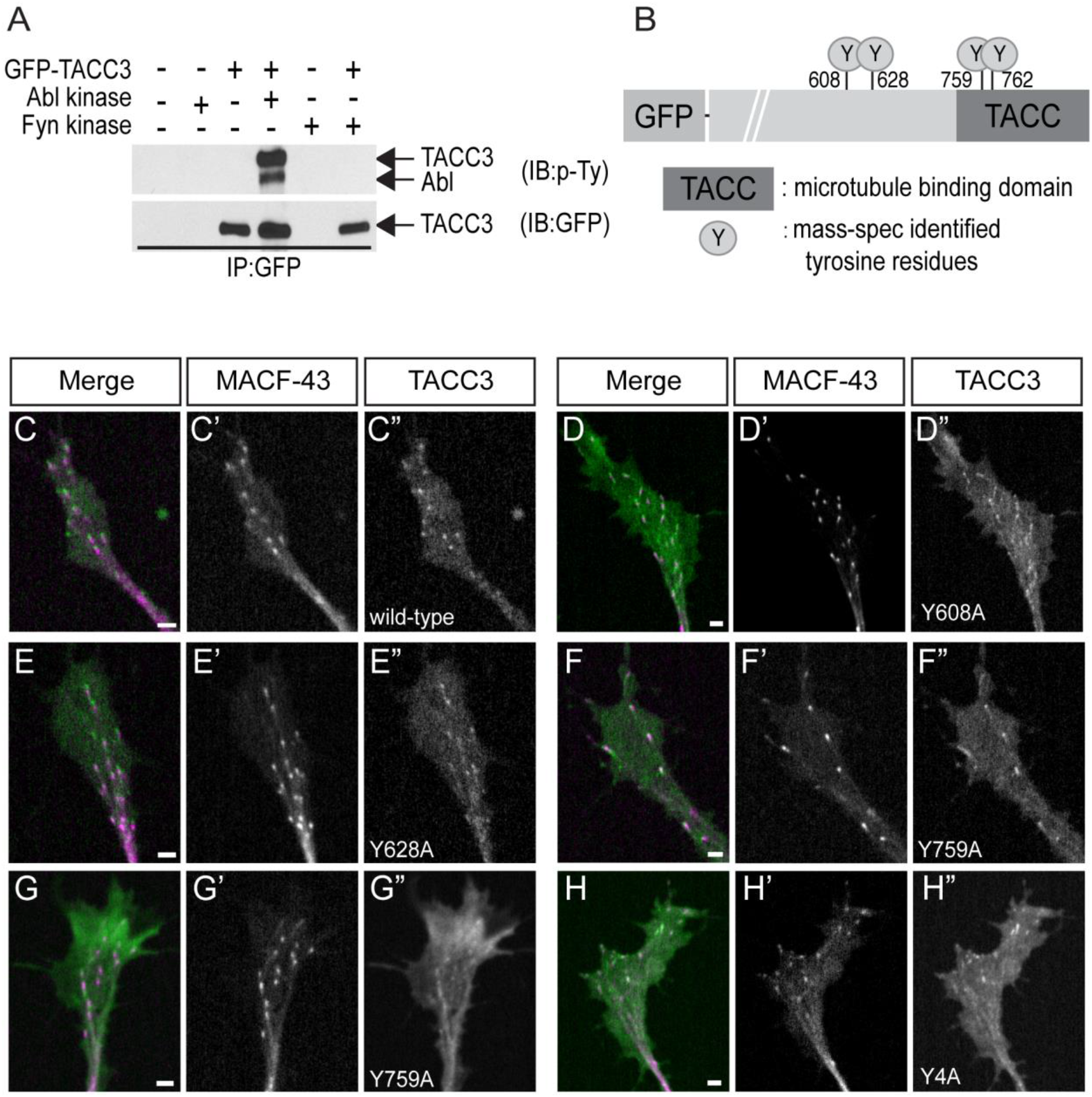
Abelson kinase induces phosphorylation of TACC3 Abelson kinase induces phosphorylation of TACC3. (A) Western blot performed with phospho-tyrosine specific antibody showing Abelson-induced tyrosine phosphorylation of full-length TACC3 (lane 4). Phosphorylation signal is not present when TACC3 is expressed alone (lane 2) or with another tyrosine kinase Fyn (lane 6). (B) Cartoon showing mass-spec-identified tyrosine residues of TACC3 targeted by Abl phosphorylation. (C-H) Confocal images of growth cones expressing MACF-43 (magenta) and TACC3 (green) with phospho-null mutations at identified residues, showing localization of TACC3 phospho-null mutants to microtubule plus-ends. Scale bar, 2μm.

To study the role of TACC3 phosphorylation, we next wanted to identify the tyrosine residues that are targeted by Abl. Mass-spec analysis of full-length TACC3 identified 4 tyrosine residues: two (Y759, Y762) in the conserved TACC domain, the domain that is responsible for microtubule plus-end binding, and two (Y608, Y628) outside of the TACC domain (Fig. 1B). These phosphorylation events only occurred when Abl was co-expressed with TACC3. To assess the importance of these tyrosine residues, we generated single and combinatorial phospho-null mutants of TACC3 by substituting tyrosine with alanine or phenylalanine. However, neither single nor combinatorial phospho-null mutations caused a reduction in tyrosine phosphorylation levels, determined by Western blot analysis (Supplementary Figure 1A-D). Moreover, all of the single and combinatorial mutants were still able to track microtubule plus-ends (Fig. 1C-H, Supplemental Movies 1-6). This initial examination suggests that other tyrosine residues might also be phosphorylated in TACC3 and thus contribute to the overall phospho-tyrosine signal detected by Western blot.

### Tyrosine phospho-null mutations impair the localization of the TACC domain at microtubule plus-ends in growth cones and mesenchymal cells

Since none of the TACC3 phospho-null mutants based on mass-spec-identified tyrosine residues showed a reduction in phosphorylation signal, and they were all still able to localize to microtubules, we decided to examine other tyrosine residues within TACC3. The full-length TACC3 possesses a total of 11 tyrosine residues throughout the entire protein. Six of these tyrosine residues (Y725, Y759, Y762, Y832, Y846, Y857) are in the TACC domain (aa715-aa931), one (Y130) is in the N-terminal domain (aa1-aa133), and four (Y320, Y428, Y608, Y628) are in the middle domain (aa134-aa634) (Fig. 2A).

**Figure 2.**
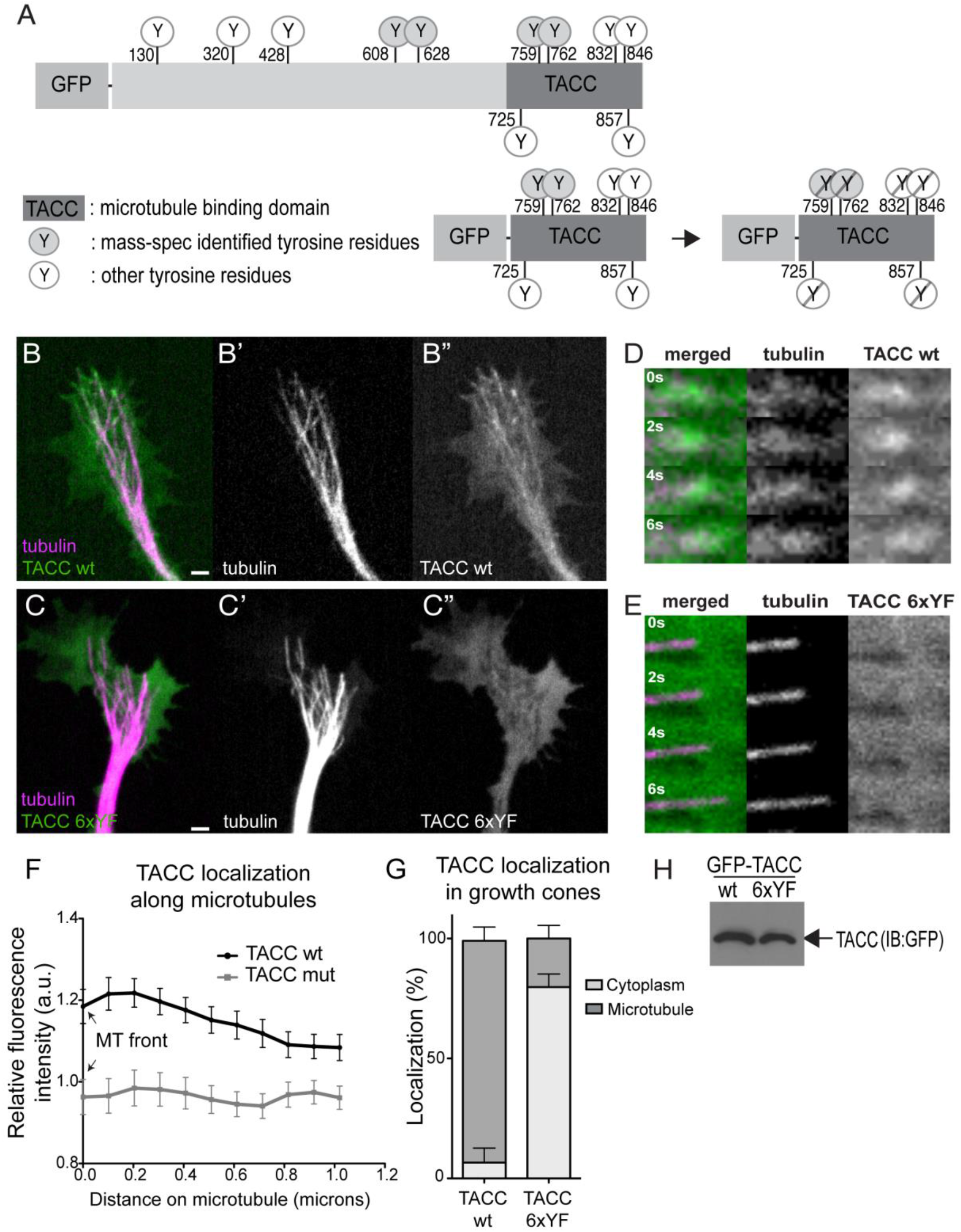
Tyrosine phospho-null mutations impairs the localization of TACC domain at microtubule plus-ends in growth cones and mesenchymal cells. (A) Cartoon showing mass-spec-identified tyrosine residues and other tyrosine residues of TACC3 (top) and the TACC domain with residues mutated to phenylalanine in the GFP-TACC phospho-null 6xYF construct. (B-C) Confocal images of neuronal growth cones, obtained from time-lapse recordings expressing tubulin (magenta) (B’, C’) and TACC (green) wild-type (B”) or tyrosine phospho-null mutant (C”). (D-E) Magnified montages of time-lapse sequences of single microtubule. TACC wild-type (green) localizes to microtubule (magenta) plus-ends (D) while TACC tyrosine phospho-null mutant (green) is absent from microtubule (magenta) plus-ends and remains mostly cytoplasmic (E). (F) Fluorescent intensity profile (y-axis) along microtubules (x-axis) determined by line-scan analysis showing green fluorescent intensity (TACC wt or TACC p-null mutant) relative to background. Line is drawn starting from the beginning of the microtubule plus-end. (G) Plot showing the quantification of microtubule plus-end versus cytoplasmic localization of TACC constructs within the growth cones as percentage of total growth cones examined. On average, TACC wild-type shows microtubule plus-end binding in 92.5% of the growth cones examined while TACC tyrosine p-null mutant showed plus-end localization in 20% of the growth cones and remained mostly cytoplasmic in 80% of the growth cones examined. (H) Western blot performed with GFP antibody showing the expression of GFP-TACC wild-type and GFP-TACC phospho-null mutant. Scale bar, 2μm.

The C-terminal TACC domain of the TACC3 protein shows high sequence similarity among vertebrates as well as among other TACC family members (Peset and Vernos, 2008). Moreover, the TACC domain is the portion of the TACC3 protein that is responsible for its microtubule plus-end tracking behavior (Nwagbara et al., 2014) and also for interacting with the microtubule polymerase XMAP215 (Mortuza et al., 2014). Therefore, we decided to specifically investigate the importance of tyrosine residues within the TACC domain to determine whether they might be contributing to the Abl-induced phosphorylation we observed earlier. To investigate this, we mutated all six of the tyrosine residues within the TACC domain to phenylalanine (Fig. 2A) and tested these mutants for their phosphorylation status. Surprisingly, despite lacking a phosphorylatable tyrosine with the TACC sequence, GFP-TACC phospho-null (p-null) mutants showed similar phospho-tyrosine signals compared to the wild-type TACC domain, as verified with Western blot (Supplementary Figure 2A-B). Since there are no tyrosines left in the TACC domain, the only other source that might contribute to the observed Western blot signal is GFP. Therefore, we performed a Western blot for the GFP tag alone with a phospho-tyrosine specific antibody, and we found that GFP was indeed getting phosphorylated specifically in the presence of Abl kinase (Supplementary Figure 2C-D). Thus, GFP itself was probably contributing to the overall signal that we obtained with the GFP-tagged TACC constructs previously. However, while Abl-induced phosphorylation of GFP muddled the Western blot results, we still had previously determined that tyrosine residues of TACC3 were indeed being phosphorylated as a result of Abl signaling, and we decided to pursue whether tyrosine phosphorylation directly impacted the ability of the TACC domain to bind microtubules.

When we examined whether TACC protein with tyrosine phospho-null mutations (TACC 6xYF) could still localize to microtubules, we found that both full-length TACC3 (data not shown) and TACC domain-only phospho-null mutants showed changes in their microtubule localization (Fig. 2B-C, Supplemental Movie 7-8). In both neuronal growth cones (and mesenchymal cells, not shown) isolated from *Xenopus* embryos, TACC phospho-null mutants showed less localization along microtubules, determined by line-scan averages of fluorescent intensities obtained from microtubule plus-ends (Fig. 2D-E, F). TACC phospho-null mutant remained cytoplasmic in 80% of the growth cones and showed microtubule localization in 20% of the growth cones examined, while wild-type TACC localized at microtubule plus-ends in 93% of growth cones examined (Fig. 2G). We also confirmed protein expression levels by Western blot and showed that the GFP-TACC mutant expression was comparable to GFP-TACC (Fig. 2H). These data suggest that retaining phosphorylatable tyrosines within the TACC domain is important for TACC localization along microtubules.

### TACC tyrosine phospho-null mutant-expressing axons grow less persistently, thereby resulting in shorter axons compared to TACC wild-type-expressing axons

We have previously shown that TACC3 binding to microtubules is critical for promoting axon outgrowth (Nwagbara et al., 2014). To test whether phosphorylation of the TACC domain is important in regulating axon outgrowth parameters, we measured the length of axons in cultured *Xenopus laevis* neural tube explants expressing either TACC wild-type (wt) protein or TACC phospho-null mutant protein. While expression of TACC wt led to formation of longer axons, expression of the TACC phospho-null mutant (0.86 ± 0.04 N=155) was unable to promote increased axon outgrowth, and instead, showed a 17% reduction in axon length compared to controls (1.00 ± 0.03 N=277) and a 34% reduction compared to TACC wt (1.15± 0.06 N=199) (Fig. 3A).

**Figure 3.**
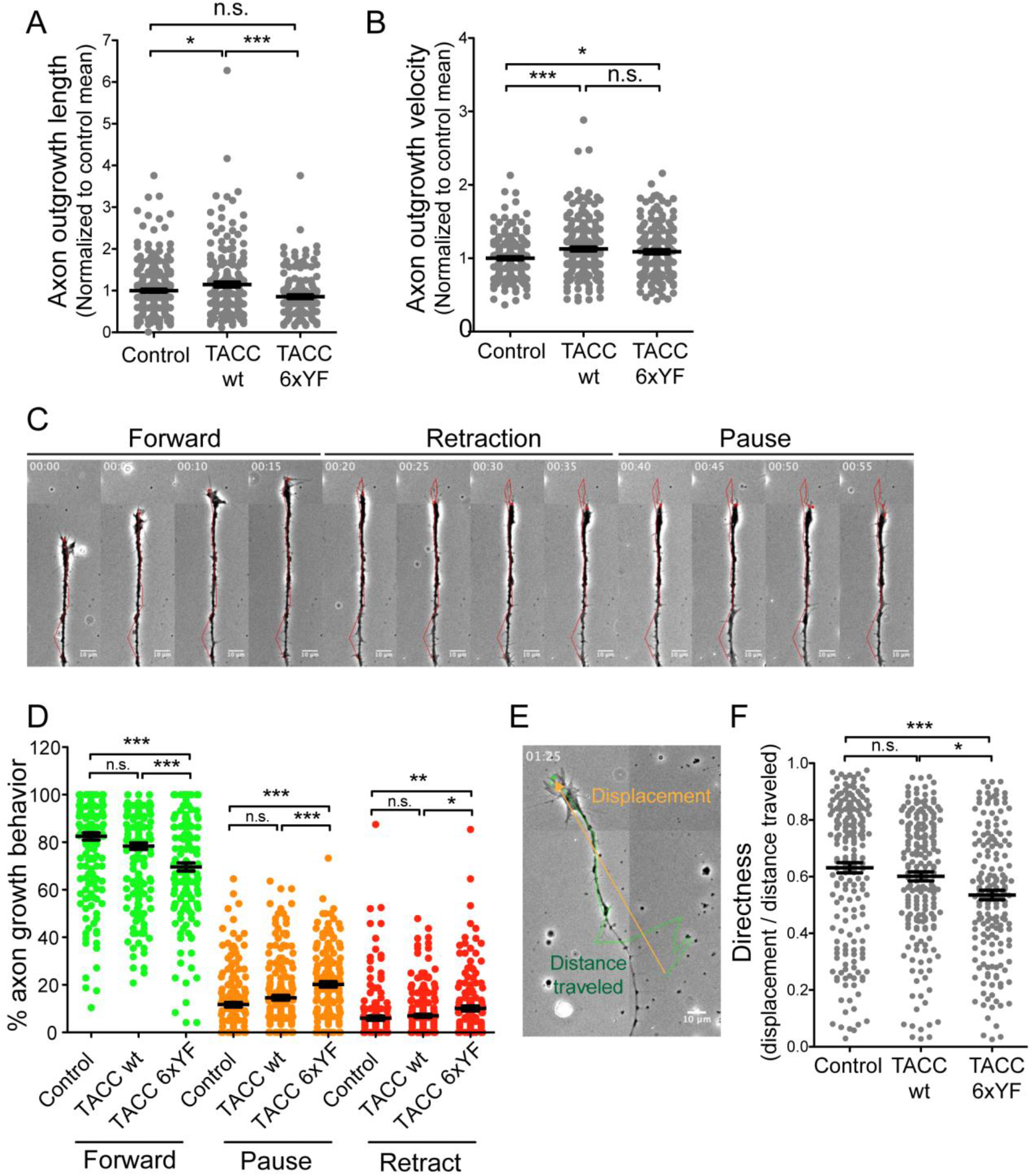
TACC tyrosine phospho-null mutant expressing axons grow less persistently thereby grow shorter axons compared to TACC wt axons. (A) Quantification of axon length in cultured neural tube explants shows TACC phospho-null mutant expressing neurons grow axons shorter by 16.8% compared to control and 34% compared to TACC wt, while TACC wt expressing explants grow axons longer by 12.8% compared to control. (B) Measurement of axon forward movement velocity showing TACC wt increases axon forward movement velocity by 10% compared to control while TACC phospho-null mutant increases by 8%. (C) Representative phase contrast image montage of an axon depicting phases of forward movement, retraction and pause. (D) Plot showing the percentage of forward movement (green), pause (orange) and retraction (red) of axon growth. TACC phospho-null mutant expressing axons spend 18% less time moving forward compared to control and 12% compared to TACC wt (green dots). TACC phospho-null mutant axons (20.20 ± 1.150 N=168) spent 42% more time pausing compared to control and 27% compared to TACC wt (orange dots). They also tend to retract more frequently; 39% compared to control and 30% compared to TACC wt. (E) Representative image of an axon growth track depicting displacement and distance traveled between t=0 and t=1h 25min (F) Plot showing the directness of axon growth. TACC phospho-null mutant axons grow 12% less directly compared to TACC wt) and 17% compared to control axons. TACC wt axons also grow 5% less directly compared to control but it was not significant. The asterisk (*) indicates statistical significance with α < 0.05 * P < 0.05, ** P < 0.01, **** P < 0.0001, ns: not significant, from an ANOVA analysis comparing multiple conditions with Tukey’s post hoc analysis. Values given are mean of normalized data pooled from independent experiments.

To further assess the difference in axon length between the neurons expressing TACC wt and the phospho-null mutant, we measured axon outgrowth velocities. Interestingly, despite the fact that phospho-null mutant-expressing axons are not as long as TACC wt-expressing axons, we found only minor (insignificant) differences in outgrowth velocity. The average normalized axon forward movement velocity was 8% faster in TACC phospho-null mutant conditions (1.10± 0.03 N=172, **p=0.0082) compared to controls (1.00 ± 0.02 N=198) and only 4% slower, compared to TACC wt (1.13 ± 0.03 N=217, not significant p=0.3). Average outgrowth velocity in TACC wt-expressing axons, on the other hand was 10% faster compared to control axons (Fig. 3B). Our data suggests that TACC phospho-null mutant-expressing axons grow at similar rates to the TACC wt-expressing axons, suggesting that the reduced axon length in the phospho-null condition could be due to another parameter of axon growth other than outgrowth velocity.

To further investigate this, we also tracked other growth behaviors of axons and recorded the number of frames that they moved forward, paused and/or retracted over the course of 4 hour long time-lapse imaging (Fig 3C, Supplemental Movie 9-10, intervals were every 5 minutes). We found that TACC phospho-null mutant-expressing axons (69.61 ± 1.77 N=168) spent 18% less time moving forward compared to controls (82.30 ± 1.5 N=183, ***p<0.0001) and 12% less compared to TACC wt (78.20 ± 1.43 N=218, ***p<0.0001) (Fig 3D, green dots). Additionally, TACC phospho-null mutant axons (20.20 ± 1.150 N=168) spent 42% more time pausing compared to controls (11.67 ± 1.00 N=183, ***p<0.0001) and 27% more compared to TACC wt 14.67 ± 0.97 N=218, ***p=0.0003) (Fig 4D, orange dots). They also tended to retract more frequently: 39% compared to controls (6.125 ± 0.89 N=183, **p=0.0043) and 30% compared to TACC wt (7.098 ± 0.67 N=218, *p=0.0132) (Fig 3D, red dots, Supplemental Movie 11).

**Figure 4.**
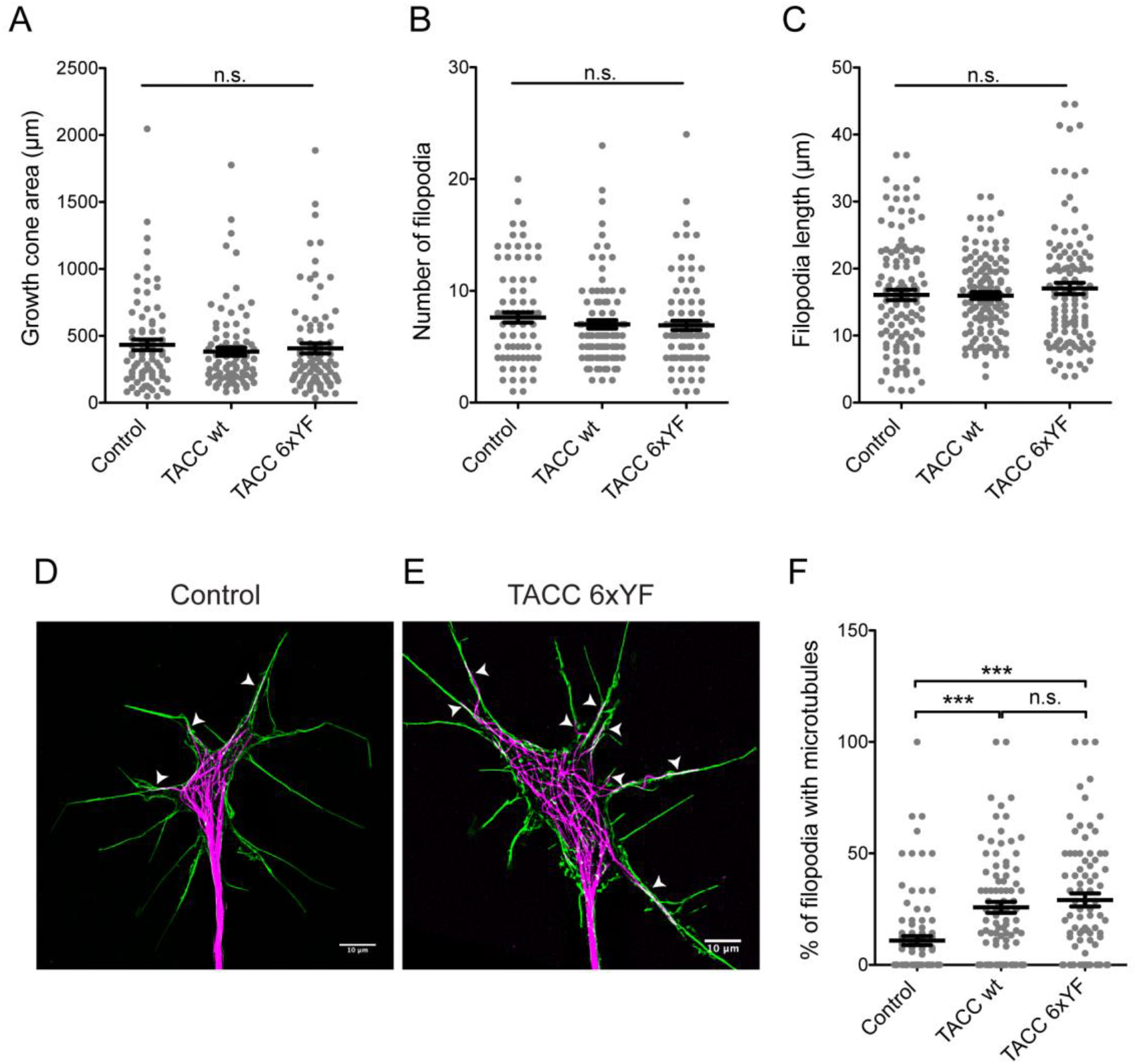
TACC tyrosine phospho-null mutant-expressing growth cones display increased numbers of filopodia that contain microtubules. (A-C) The average growth cone area of TACC mutant (406.5 ± 37.17 μm, N=88) expressing growth cones was 13% smaller than control (432.6 ± 39.62 μm, N=77) and 6% larger than TACC wt (382.5 ± 30.17 μm, N=94) (A). TACC p-null mutant growth cones had an average of 6.9 ± 0.4130 N=93 filopodia which was 10% less than control growth cones (7.6 ± 0.4510 N=89) while TACC wt had 9% fewer filopodia than control (7.0 ± 0.3812 N=101) (B). The average filopodia length in TACC p-null mutant growth cones was 17.03 ± 0.8448μm, N=113, while it was 16.07 ± 0.7707 μm, N=119 in control growth cones, and 15.96 ± 0.5079 μm, N=126 in TACC wt expressing growth cones (C). Almost 30% of the filopodia had microtubules in the TACC p-null mutant growth cones (29.13 ± 2.932 percent of total filopodia examined, N=82, E), which is 63% more than control growth cones (10.92 ± 2.011 percent of total filopodia examined, N=90, D), that had only 11% of their filopodia invaded by microtubules. TACC wt-expressing growth cones also had a greater number of filopodia with microtubules (25.82 ± 2.446 N=92) compared to controls. While not significant, TACC p-null growth cones had 11% more filopodia with microtubules compared to TACC wt (F). Scale bar 10μm.

In addition to pause and frequency rates, we also examined the directness of axon outgrowth. The still images of neural tube explants that were used to measure axon length show the final displacement of an axon. However, axons do not necessarily follow a linear trajectory as they grow. In fact, they often spend time wandering, which is part of their exploratory behavior. Therefore, the directness of outgrowth can be determined by dividing the displacement distance by the total distance traveled (Fig. 3E). We found that TACC phospho-null mutant expressing-axons (0.5353 ± 0.02 N=179) grow 12% less directly compared to TACC wt-expressing axons (0.6012 ± 0.02 N=195) and 17% less compared to control axons (0.6313 ± 0.02 N=192, Fig. 3F). These data suggest that while axon outgrowth speed is not affected by TACC tyrosine phospho-null mutations, growth persistency, which is determined by pause and retraction frequency, as well as growth directionality, both seem to be impaired in TACC phospho-null mutant expressing axons. Thus, this could explain the shorter axon length that we observed in TACC phospho-null mutant-expressing neural tube explants.

### TACC tyrosine phospho-null mutant-expressing growth cones display increased numbers of filopodia that contain microtubules

Axons of neurons expressing the TACC phospho-null mutant tend to stop and retract more frequently compared to those expressing wild-type TACC. Additionally, we observe that their growth is less directed compared to TACC wt and controls. An inverse correlation between growth cone advance rate and growth cone size has been reported previously (Ren and Suter, 2016). Therefore, we became interested in examining the growth cone size, along with other morphological features, to see whether increased pause and retraction rates in TACC phospho-null mutant-expressing growth cones might also occur alongside changes in growth cone morphology.

We initially examined the growth cone area, filopodia length, and filopodia number in cultured neural tube explants that are fixed and stained for microtubules (tubulin) and actin (phalloidin), followed by high resolution Structured Illumination Microscopy (SIM) imaging. Somewhat surprisingly, we found no significant difference in any of the morphological features examined (Fig. 4A-C). The average growth cone area of TACC phospho-null mutant-expressing growth cones (406.5 ± 37.17 μm, N=88, p=0.6) was 13% smaller than control (432.6 ± 39.62 μm, N=77) and 6% larger than TACC wt (382.5 ± 30.17 μm, N=94, p=0.6) (Fig. 4A). The average filopodia number did not differ across conditions either (p=0.4). TACC phospho-null mutant growth cones had an average of 6.9 ± 0.41 N=93 filopodia, which was 10% less than control growth cones (7.6 ± 0.45 N=89) while TACC wt had 9% fewer filopodia than control (7.0 ± 0.38 N=101) (Fig. 4B). The average filopodia length in TACC phospho-null mutant growth cones was 17.03 ± 0.84μm, N=113, while it was 16.07 ± 0.77 μm, N=119 in control growth cones, and 15.96 ± 0.51 μm, N=126 in TACC wt expressing growth cones (Fig. 4C).

However, while average filopodia number was similar among conditions, the number of filopodia that contained microtubules was found to be significantly higher in the TACC phospho-null mutant-expressing growth cones. Almost 30% of the filopodia had microtubules in the TACC phospho-null mutant growth cones (29.13 ± 2.932 percent of total filopodia examined, N=82), which is 63% more than control growth cones (10.92 ± 2.011 percent of total filopodia examined, N=90, ***p<0.0001), that had only 11% of their filopodia invaded by microtubules. TACC wt-expressing growth cones also had a greater number of filopodia with microtubules (25.82 ± 2.446 N=92, ***p<0.0001) compared to controls. While not significant, TACC phospho-null growth cones had 11% more filopodia with microtubules compared to TACC wt (Fig. 4F). Thus, it appears that over-expression of the phospho-null TACC domain somehow leads to an increased number of microtubules that penetrate into filopodia, which may directly or indirectly contribute to increased pausing and retraction rates of these axons.

### TACC tyrosine phospho-null mutant-expressing axons are more responsive to repellent guidance signals

We previously tested TACC3 function downstream of Slit2 and found that growth cones overexpressing TACC3 were more resilient to bath-applied Slit2-induced growth cone collapse compared to control growth cones (Erdogan et al., 2017). While bath application of guidance proteins is useful to test how manipulation of protein levels would alter the growth cone’s reaction to an applied guidance molecule, it does not convey information regarding the ability of the growth cone to make guidance choices such as steering. In order to test that, we utilized an approach where the repellent guidance protein Ephrin-A5 is coated on a glass coverslip in zigzag patterns and neural tube explants are cultured on top of these cue-coated coverslips. After 12-18 h of culturing, explants are fixed and stained for tubulin and actin to examine axon responsiveness to Ephrin-A5. The responsiveness is scored based on how many of the axons that grow out from the given explant preferred to stay on the Ephrin-A5-free (growth permissive) surface versus how many of them cross that barrier and grow on the Ephrin-A5 (non-permissive) surface.

We found that control neural tube explants (Fig. 5A), composed of a heterogeneous population of neurons, do not show a preference towards a particular surface and grow axons on both permissive (Ephrin-A5 free) and non-permissive (Ephrin-A5) surfaces equally (Fig. 5D; Control - off, 50.01 ± 2.93 N=55; Control- on, 49.99 ± 2.93, respectively, p=0.9). However, TACC wt over-expressing neural tube explants (Fig. 5B) show a preference for the permissive surfaces, as they grow 25% more axons on permissive surfaces (Fig. 5D; 57.18 ± 2.68 N=62, TACC wt – off, ***p=0.0002) compared to the Ephrin-A5 coated surface (TACC wt - on, 42.82 ± 2.68 N=62). TACC phospho-null mutant-expressing axons (Fig. 5C), on the other hand, show an even stronger preference for the permissive surface, as they grow 41.3% more axons on the permissive surface (Fig. 5D; TACC mut - off, 63.02 ± 3.03 N=42, ***p<0.0001) compared to the Ephrin-A5 coated non-permissive surface (TACC mut - on, 36.98 ± 3.03 N=42). Together, these data suggest that expression of the TACC domain with the non-phosphorylatable tyrosine residues are less able to grow on the Ephrin surface and are thus more responsive to the Ephrin-A5 repellent guidance cue activity.

**Figure 5.**
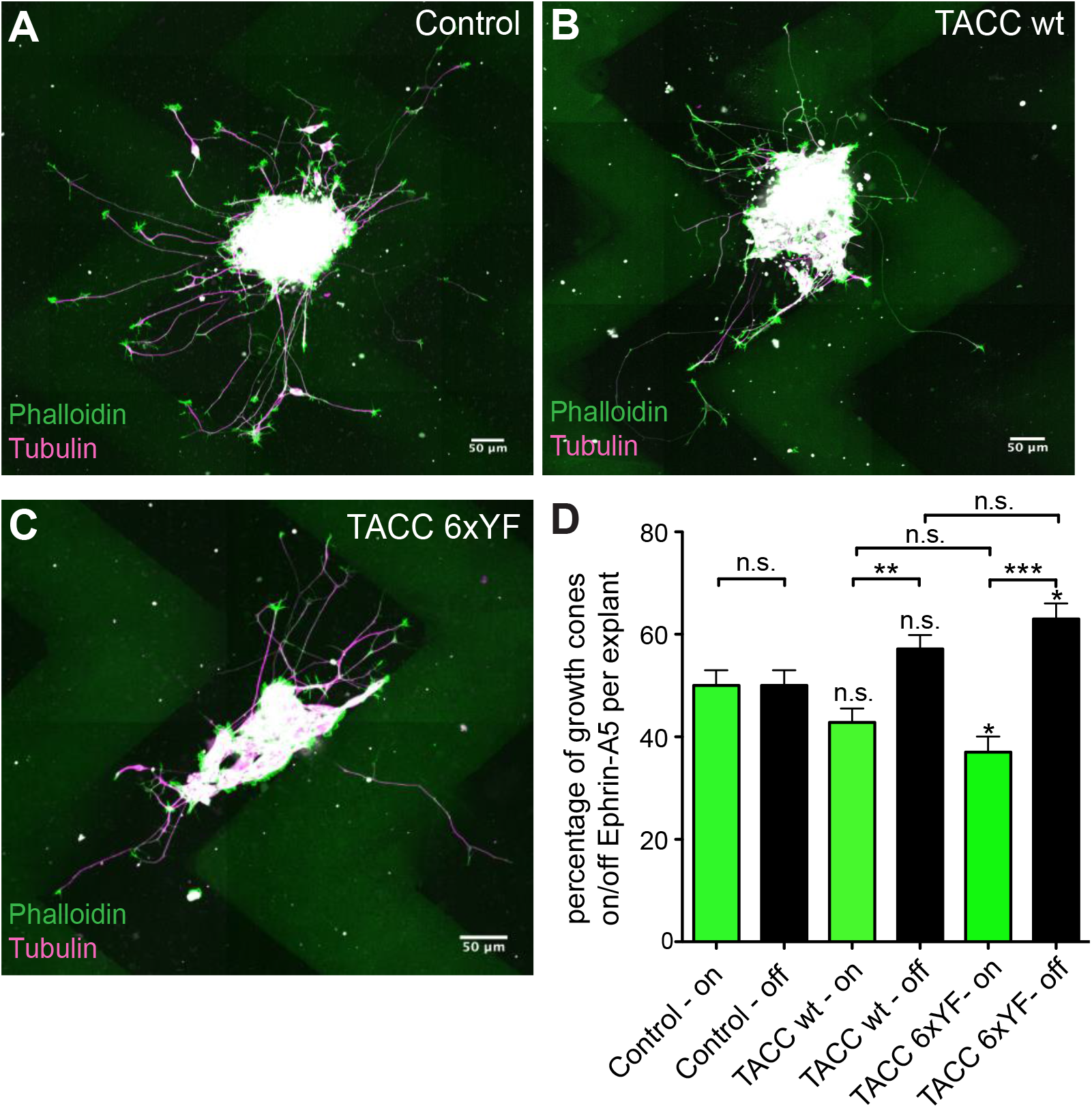
TACC tyrosine phospho-null mutant-expressing axons are more responsive to repellent guidance signals. (A-C) Immunofluorescence images of control (A). TACC wt (B) and TACC p-null mut (C) labelled with phalloidin (green) to label growth cones and tubulin (red) to label axons showing axon responsiveness to Ephrin-A5 coated zigzag surfaces (green). (D) Quantification of the number of axons on permissive (Ephrin-A5 free) versus non-permissive (Ephrin-A5) surfaces. Control neural tube explants does not show a preference between permissive versus non-permissive surfaces and grow axons on both surfaces equally (Control - off, 50.01 ± 2.930 N=55; Control- on, 49.99 ± 2.930, respectively). TACC wt expressing neural tube explants grow 25% more axons on permissive surfaces (57.18 ± 2.683 N=62, TACC wt - off) compared to Ephrin-A5 coated surface (TACC wt - on, 42.82 ± 2.683 N=62). TACC p-null mutants grow 41.3% more axons on permissive surface (TACC mut - off,63.02 ± 3.036 N=42) compared to Ephrin-A5 coated non-permissive surface (TACC mut - on, 36.98 ± 3.036 N=42). Scale bar 50μm.

## Discussion

### Abelson kinase induces phosphorylation of TACC3

In this study, we sought to examine the impact of TACC3 tyrosine phosphorylation on its interaction with microtubules and regulation of axon outgrowth and guidance. We showed that co-expressing TACC3 with Abelson kinase in HEK293 cells induces TACC3 tyrosine phosphorylation, as evident by the phospho-tyrosine signal we obtained by Western blot. Our Western blot analysis also shows that phosphorylation of TACC3 happens only in the presence of Abelson, as we do not observe any phospho-tyrosine signal when TACC3 is expressed alone or with Fyn, which is another tyrosine kinase (Fig. 1A).

Interestingly, creating single or combinatorial phospho-null mutations at tyrosine residues that we identified as potential targets for Abelson did not show any changes in phospho-tyrosine signal levels observed with Western blot. In an attempt to identify the source of tyrosine phosphorylation, we tested other tyrosine residues and focused specifically on the ones in the TACC domain positioned at the C-terminal of TACC3 and which is responsible for microtubule plus-end tracking and interaction with its well-studied partner, microtubule polymerase XMAP215. The TACC domain possesses 6 tyrosine residues with two of them (Y759 and Y762, aa location given based on full length protein) identified as Abelson targets from the mass-spec analysis and three of them (Y832, Y846 and Y857) predicted as putative targets for Abl via *in silico* analysis. Intriguingly, mutating all these tyrosines into non-phosphorylatable phenylalanine did not cause any reduction in the phospho-tyrosine signal level as evident by the Western blot (Supplementary Fig. 2A-B). In spite of the lack of any tyrosines in the TACC domain, it was intriguing to see that there was still a phospho-tyrosine signal in the Western blot. The only possible source of phosphorylatable tyrosine was GFP that is tagged to the TACC domain. Tyrosine phosphorylation of GFP has not been reported previously, to our knowledge, nor does *in silico* phosphorylation prediction identify any sites by which kinases might target GFP. However, expression of GFP with Abelson clearly showed a strong phospho-tyrosine signal (Supplementary Fig. 2C-D) suggesting that GFP might be partially contributing to the phospho-tyrosine signal that we have been observing in GFP-TACC.

### Tyrosine residues within the TACC domain are important for TACC localization to microtubules

The impact of phosphorylation on the interaction between +TIPs and microtubules has been studied for several +TIPs, such as CLASP (Hur et al., 2011; Kumar et al., 2012; Kumar et al., 2009; Watanabe et al., 2009), APC (Zhou et al., 2004), ACF7 (Wu et al., 2011), EB1 (Zhang et al., 2016). Here, we demonstrated that tyrosine residues within the TACC domain of TACC3 are important for maintaining the interaction between TACC3 and microtubules. The TACC domain with all tyrosine residues mutated into phenylalanine remains mostly cytoplasmic, while TACC wild-type localizes to microtubules. It should be noted that the wild-type TACC domain, in contrast to full-length TACC3 (which shows primarily plus-end binding), also shows lattice binding in addition to plus-end localization. Consistent with our previous observations (Nwagbara et al., 2014), this suggest that N-terminus of TACC3 is important for restricting TACC3 localization to the microtubule plus-ends.

+TIPs track microtubule plus-ends either autonomously, through recognition of the growing microtubule structure, or non-autonomously, through an interaction with another plus-end tracking protein such as end-binding (EB) proteins. Although the mechanism of how TACC3 tracks plus-ends is not fully resolved, EB-dependent plus-end tracking can be ruled out as TACC3 does not contain a SxIP motif (serine-any amino acid-isoleucine-proline) that is required for EB binding. It is believed that TACC3 tracks microtubule plus-ends through its interaction with XMAP215, which is mediated by the TACC domain (Kinoshita et al., 2005; Mortuza et al., 2014; Peset et al., 2005). Therefore, it is possible that the impaired interaction between microtubules and TACC phospho-null mutant could be arising due to a change in TACC3’s ability to interact with XMAP215.

The TACC domain consists of two coiled-coil domains, which means that the sequence follows heptad repeats. Presence of these coiled-coil domains are responsible for TACC3’s oligomerization, which is important for TACC3’s function and its interaction with XMAP215 (Mortuza et al., 2014; Thakur et al., 2014). Using Multicoil, a coiled-coil prediction program, we looked at whether tyrosine-to-phenylalanine mutations might have altered coiled-coil formation or dimerization of TACC domain. Based on this prediction, it seems as though the TACC phospho-null mutant is still likely able to dimerize and form a coiled-coil (Supplementary Figure 3), suggesting that the interaction between XMAP215 and TACC phospho-null mutant might be retained.

Although *in silico* analysis suggests that switching from phosphorylatable tyrosine to non-phosphorylatable phenylalanine does not impact TACC3’s coiled-coil structure, reduced interaction between phospho-null TACC mutants and microtubules might be explained by a change in electrostatic interactions between the two. While this could be a possibility, negative charges introduced via phosphorylation often cause dissociation of proteins from the microtubule (Hur et al., 2011; Iimori et al., 2012) which is already loaded with negative charges due to negatively-charged residues at the C-terminus of tubulin. While our findings suggest that the tyrosine residues of TACC domain are important for mediating TACC3’s interaction with microtubules, the exact mechanism of how they are involved in this interaction remains to be determined.

### TACC phospho-null mutant axons are shorter due to increased pausing and retraction

Microtubule advance within the growth cone is important for consolidating and driving growth cone forward movement (Lowery and Van Vactor, 2009). Intriguingly, despite increasing microtubule growth speed and length, our data shows that TACC phospho-null mutant-expressing axons of cultured neural tube explants are shorter in spite of slightly higher axon forward velocity rates compared to control. When we further examined other axon outgrowth parameters, we found that TACC phospho-null mutants tend to pause and retract more frequently compared to TACC wild-type and controls (Fig. 3D). Additionally, mutant axons tend to grow with less directionality (Fig. 3F).

In an attempt to seek further explanation to axon outgrowth behavior, we examined growth cone morphology, since growth cone morphology can often predict the growth behavior of axon (Ren and Suter, 2016). For example, increased growth cone size is often associated with frequently pausing or slow growing axons. Although we did not see a significant difference in growth cone size, filopodia length, or number among conditions (Fig. 4A-C), there was a significant increase in the number of filopodia with microtubules in the TACC phospho-null mutant-expressing growth cones (Fig. 4F).

We know that microtubules can play an instructive role in growth cone directional motility. Having more filopodia with microtubules might generate more alternative routes for the growth cone to extend, which could result in more pausing for decision making and could also cause the growth cone to wander more, which would result in less directed movement. Additionally, there might be increased microtubule/F-actin coupling which would enable microtubules to track more persistently along F-actin into the filopodia. In fact, increased microtubule/F-actin coupling could also explain increased retraction rates. For example, actin filaments in the growth cone are subjected to rearward translocation also known as retrograde flow, due to myosin II activity (Suter and Forscher, 2000). The retrograde flow of actin filaments can be attenuated when the F-actin cytoskeleton engages with receptors at point contact sites. This interaction acts as a molecular clutch that will restrain actin retrograde flow, while continuing actin polymerization will generate the force to allow membrane protrusion, thereby growth cone advance. The growth cone retracts when actin retrograde flow fails to be attenuated, which is also an indication of poor surface adhesion. It is known that dynamic microtubules play an important role in facilitating focal adhesion dynamics by transporting molecules that are involved in focal adhesion turnover (Stehbens and Wittmann, 2012). Several microtubule plus-end binding proteins have been studied for their association with adhesion sites (Stehbens et al., 2014; Zhang et al., 2016). Therefore, by facilitating microtubule protrusion into the filopodia, the TACC phospho-null mutant domain might be playing an indirect a role in adhesion dynamics.

A potential role in point contact regulation can also explain avoidance of EphrinA5-coated repellent substrates. Filopodia with microtubules could be more sensitive to cues, due to the microtubule-mediated signal trafficking, thereby affecting the interaction with the underlying substrate. Moreover, as speculated earlier, having more filopodia with microtubules could generate more alternative routes; thus, when the growth cone encounters a non-permissive substrate, it could pick one of these alternative routes and steer away from the repellent source.

In conclusion, we have demonstrated that tyrosine residues within the TACC domain, which is the domain important for mediating microtubule plus-end tracking behavior, are critical for localizing the protein to microtubules, regulating axon outgrowth parameters and making guidance decisions. Given the increased number of microtubules in the filopodia when the TACC domain is over-expressed, we hypothesize that the tyrosine residues in the TACC domain might be involved in regulating a potential TACC-mediated interaction between microtubules and F-actin. Such an interaction has not been proposed for TACC3 before, however, there are studies that might offer a potential interaction between TACC3 and the actin cytoskeleton. A proteomic screen previously identified an interaction between TACC3 and actin regulator proteins Ena/VASP (Hubner et al., 2010). Additionally, the close interactor of TACC3, XMAP215, has recently been shown to interact with actin and mediate microtubule/F-actin interaction in growth cones (Slater et al., 2019), which makes TACC3 a candidate to be involved in actin cytoskeleton interaction either direct or indirectly.

To our knowledge, TACC3 tyrosine phosphorylation has not been explored extensively before. Nelson *et. al.* identified two tyrosine residues, Y684 and Y753 (corresponding to residues Y725 and Y832 in *X. laevis* and conserved across species, Supp. Fig. 4) that showed enhanced phosphorylation when it is fused to Fibroblast Growth Factor Receptor 3 (FGFR3) (Nelson et al., 2016). It has been indicated that the FGFR3–TACC3 fusion is important for the activation of FGFR3 tyrosine kinase activity, and fusion of these proteins increases cell proliferation and tumor formation. Moreover, TACC3 in association with ch-TOG has been shown to localize to the mitotic spindles (Lee et al., 2001). However, the FGFR3-TACC3 fusion, which pulls TACC3 away from mitotic spindles and localizes to the spindle poles, causes mitotic defect,s which might explain the involvement of these proteins in tumor formation (Sarkar et al., 2017). From this perspective, it might be intriguing to investigate TACC domain tyrosine residues and TACC association with microtubules during cell division.

Finally, in addition to tyrosine phosphorylation, serine/threonine phosphorylation sites within TACC3 would be worthwhile to explore, given that there are several S/T kinases that operate under guidance cues, such as GSK3, which is a well-studied S/T kinase that is already shown to phosphorylate various +TIPs and modulate microtubule dynamics. Future work can further explore whether TACC3 is targeted by S/T kinases and whether S/T phosphorylation would affect TACC3’s function in a similar way, which could shed light on differential and asymmetric regulation of microtubule dynamics under various guidance signals.

## Materials and Methods

### Embryos

Eggs collected from *Xenopus laevis* were fertilized *in vitro* and kept between at 13-22°C in 0.1X Marc’s Modified Ringer’s (MMR). All experiments were approved by the Boston College Institutional Animal Care and Use Committee and were performed according to national regulatory standards.

### Culture of embryonic explants

Neural tubes of embryos staged according to Nieuwkoop and Faber were dissected at stages between 20-21 as described previously (Lowery et al., 2012). Neural tube explants were cultured on MatTek glass bottom dishes coated with poly-L-lysine (100 μg/ml) and laminin (20 μg/ml). Culture media prepared by mixing L-15 Leibovitz medium and Ringer’s solution supplemented with antibiotics NT3 and BDNF to promote neurite outgrowth.

### Constructs and RNA

Capped mRNAs were transcribed and purified as previously described (Lowery et al., 2013; Nwagbara et al., 2014). TACC3 pET30a was a gift from the Richter lab, University of Massachusetts Medical School, Worcester, MA and sub-cloned into GFP pCS2+. Tyrosine phospho-null mutations were introduced into wild-type GFP-TACC3 by using overlapping extension PCR method with appropriate primers designed to substitute tyrosine residues with phenylalanine to generate GFP-TACC3 6xYF. Wild-type and phospho-null mutants of TACC domain were cloned from their full-length counterparts. MACF 43 (a gift from Hoogenraad Lab) was sub-cloned into mKate2 pCS2+. Embryos either at the 2-cell or 4-cell stage were injected with the following total mRNA amount per embryo; 100 pg of GFP-MACF43 as a control for GFP-TACC3 and to analyze microtubule dynamics parameters. GFP-TACC3 full-length wild-type or phospho-null mutant injected at 1000 pg. GFP-TACC wild-type or phospho-null mutant were injected at 400pg. The human c-Abl construct was originally constructed in the Kufe lab (Harvard Medical School) (Cao et al. 2003) and gifted by Dr. Alan Howe (University of Vermont). The Fyn construct in pRK5-Entry (Mariotti et al. 2001) was acquired from AddGene (Cambridge, MA, USA).

### Cell Culture and Transfections

HEK293 cells were maintained at 37 °C and 5% CO_2_ and cultured in DMEM containing L-glutamine, sodium pyruvate and 4.5 g/L glucose (MediaTech/Corning Life Sciences, Tewksbury, MA, USA). DMEM was supplemented with 5% fetal bovine serum (Hyclone, Logan, UT, USA), 5% Cosmic calf serum (Hyclone), 50 units/mL penicillin and 50 μg/mL streptomycin (Penicillin-Streptomycin, Invitrogen, Carlsbad, CA, USA). Cells were cultured to 75% of confluence and transfected with 3-6 μg GFP-TACC3 full-length (FL) with or without 2 μg c-Abl or 3.5 μg Fyn expression plasmids using calcium phosphate precipitation. The following expression constructs encoding TACC3 phospho-null and truncation mutants were co-transfected at 3 μg with c-Abl: GFP-TACC3 Y2A (Y608A, Y628A); GFP-TACC3 Y3A (Y608A, Y628A, Y759A); GFP-TACC3 Y4A (Y608A, Y628A, Y759A, Y762A). GFP-TACC domain widtype; GFP-TACC domain phospho-null mutant (Y725F, Y759F, Y762F, Y832F, Y846F, Y857F).

### Cell Lysis, Immunoprecipitation and Western Blotting

HEK293 cells were lysed as previously described (St. Clair et al. 2019) in lysis buffer (25 mM Tris pH 7.4, 137 mM NaCl, 10% glycerol, 1% Igepal) containing protease inhibitors (5 μg/mL Pepstatin, 10 μg/mL Leupeptin, 1 mM PMSF) and phosphatase inhibitors (1 mM NaVO_3_, 25 mM NaF, 10 mM Na_2_H_2_P_2_O_7_). For cell extract immunoblotting, 20-30 μg total protein extract was denatured in protein sample buffer (125 mM Tris pH 6.8, 7.5% glycerol, 2% sodium dodecyl sulfate (SDS), 5% β-mercaptoethanol, and 0.02% bromophenol blue) at 95 °C for 5 minutes and subjected to SDS-PAGE. For immunoprecipitation experiments, 1000 μg of total protein was incubated overnight at 4 °C with 2 μg α-GFP (Life Technologies/ThermoFisher Scientific, Carlsbad, CA, USA) and 20 μL of a 50% slurry of sepharose Protein A (Rockland Immunochemicals, Limerick, PA, USA) and Protein G resin (G-Biosciences, St. Louis, MO, USA) prewashed with lysis buffer. Immune complexes were washed 3 times with lysis buffer, dried and denatured in protein sample buffer at 95 °C for 5 minutes. Denatured cell extract and immunoprecipitation samples were subjected to SDS-PAGE separation. For mass spectrometry analysis, the SDS-PAGE gel was stained with Coomassie (0.1% Coomassie brilliant blue R-250, 20% glacial acetic acid and 40% methanol) and subsequently prepared for mass spectrometry as described below. For immunoblotting experiments, the following primary antibodies were diluted in 10 mL of 1.5% BSA in TBST containing 0.0005% sodium azide and incubated at 4 °C with the membranes overnight: α-GFP (1:2000, rabbit pAb, Life Technologies); α-Abl (1:1000, rabbit pAb, Santa Cruz Biotechnology, Dallas, TX, USA); α-phosphotyrosine 4G10 (1:1000, mouse mAb, EMD Millipore, Billerica, MA, USA). The following secondary antibodies were used: α-mouse-HRP (goat IgG, 1:5000; EMD Millipore); α-rabbit-HRP (goat IgG, 1:15,000, EMD Millipore); or for immunoprecipitation experiments, α-rabbit-HRP Light Chain Specific (goat IgG, 1:10,000, Jackson ImmunoResearch Laboratories, West Grove, PA, USA). Proteins were detected using enhanced chemiluminescence (ThermoFisher Scientific, Waltham, MA, USA) and film was developed using a Medical Film Processor SRX-101A (Konica Minolta Medical and Graphic, Tokyo, Japan).

### Mass Spectrometry

To identify Abl-induced phosphorylation sites on TACC3, GFP-TACC3 was transfected with or without c-Abl. GFP-TACC3 was immunoprecipitated using α-GFP and subjected to SDS-PAGE separation and Coomassie staining as described above. The region corresponding to the molecular weight of GFP-TACC3 (180 kDa) was excised and prepared for mass spectrometry as previously described (Cheerathodi et al. 2015). Briefly, gel regions were diced and proteins were subjected to in-gel digestion with 6 ng/μL trypsin (Promega, Madison, WI, USA) in 50 mM ammonium bicarbonate at 37 °C for 10-12 hours. The tryptic peptides were resuspended in 2.5% acetonitrile, 0.15% formic acid and separated via HPLC prior to MS/MS analysis on a linear ion trap-orbitrap (LTQ-Orbitrap) mass spectrometer (Thermo Electron, Waltham, MA, USA) controlled with Thermo Xcalibur 2.1 software. Peptides were eluted and electrosprayed (2.1 kV) into the mass spectrometry as previously described (St. Clair et al. 2018). The precursor scan (scan range = 360-1700 *m*/*z*, resolution = 3.0 × 10^4^) was followed by ten collision-induced dissociation (CID) tandom mass spectra. CID spectra were acquired for the top ten ions in the precursor scan.

A SEQUEST search of the mass spectrometry data was performed using the forward and reverse *Xenopus* TACC3 Uniprot sequence requiring tryptic peptides and permitting phosphorylation of serine, threonine and tyrosine (+79.9663 Da), oxidation of methionine (+15.9949 Da), and acrylamidation of cysteine (+71.0371Da). Peptides were identified with a false discovery rate of less than 1%.

### Immunocytochemistry

Embryonic explant cultures were fixed with 0.2% Glutaraldehyde as described (Challacombe et al., 1997) and labelled with primary antibody (1:1000 diluted in blocking buffer made up by 1% non-fat dry milk in calcium and magnesium free PBS) to tyrosinated tubulin [YL1/2] (rat monoclonal, ab6160, Abcam) for 45 min at room temperature which is followed by PBS washes repeated three times and 10 min of blocking. Goat anti-rat AlexaFluor568 (1:500, ab175476, Abcam Technologies) was used as a secondary to tubulin and Phalloidin 488 (1:500, Molecular Probes) was used to label actin. Both reagents are diluted in blocking solution and applied for 45 min at room temperature followed by PBS washes several times. 90% glycerol stock was used as a mounting media for imaging.

### Stripe Assay

The stripe assay is performed as described in Knoll et al 2007 (Knöll et al., 2007). 10 ug/ml of Ephrin-A5/Fc (chimera human, Sigma, E0628) and 10 ug/ml of Fc are mixed with 2.5 ug/ ml of Anti-human IgG (Fc specific) FITC (Sigma, F9512) and Anti-human IgG (Fc specific) respectively in PBS. Solutions are incubated at room temperature for 30 min to allow for oligomerization. To coat coverslips with Ephrin-A5, a coverslip is attached to a zig-zag patterned silicon matrix and Ephrin-A5 solution is injected with a micropipet through the channel in the silicon. Injected matrices are incubated at 37°C for 30 min and then rinsed with PBS several times using the same injection method. After rinsing, coverslips are detached from the matrix and placed in a culture dish and 100 ul of second stripe solution (Fc only) is applied directly on the coverslip and incubated at 37°C for 30 min. After incubation is over coverslips are rinsed with PBS and 20 ug / ml of 100ul laminin in PBS is applied on coverslips and incubated for 1 h. After laminin incubation coverslips are rinsed with PBS several times. 400 ul of culture media is applied on coverslips and neural tube explants are placed on protein coated area and cultured for 24 h prior to imaging.

### Image acquisition and analysis

To assess axon outgrowth parameters, phase contrast images of axons were collected on a Zeiss Axio Observer inverted motorized microscope with a Zeiss 20×/0.5 Plan Apo phase objective. For axon outgrowth length, snap-shots of neural tube explants were taken 12-18h post culturing. Time-lapse images were collected for 4 h with 5 min intervals and axon growth was manually tracked frame by frame using Fiji Manual Tracking plugin. Axon growth velocity information is provided by the Manual Tracking plugin but only the forward movement velocity was included in the analysis. Axon outgrowth forward movement, pause and retraction frequencies (as a percentage of total frames tracked) are scored manually tracking axons frame by frame. High-resolution images of cultured spinal cord explants were obtained with a CSU-X1M 5000 spinning-disk confocal (Yokogawa) on a Zeiss Axio Observer inverted motorized microscope with a Zeiss Plan-Apochromat 63×/1.40 numerical aperture lens. Images were acquired with an ORCA R2 charge-coupled device camera (Hamamatsu) controlled with Zen software. For microtubule dynamics and TACC localization experiments, images are time lapse images are acquired for 1 min with 2 s intervals.

Structured illumination super-resolution images were collected on a Zeiss Axio Observer.Z1 for super-resolution microscope with Elyra S.1 system, utilizing an Objective Plan-Apochromat 63×/1.40 oil (DIC). Images were acquired with a PCO-Tech Inc.pco.edge 4.2 sCMOS camera. The images were obtained in a chamber at approximately 28°C and utilizing the immersion oil Immersol 518F 30°. Channel alignment and structured illumination processing were applied to the super-resolution images using the Zeiss Black program. Experiments were performed multiple times to ensure reproducibility. Graphs were made in GraphPad Prism. Statistical differences were determined using unpaired two tailed t-tests when comparing two conditions and one-way analysis of variance (ANOVA) with Tukey’s *post-hoc* analysis when multiple conditions were compared.

## Acknowledgements

We thank members of the Lowery Lab for helpful discussions, suggestions, and editing, especially Sangmook Lee, Paula Slater, Beth Bearce and Micaela Lasser. We thank Nancy McGilloway and Todd Gaines for excellent *Xenopus* husbandry. We also thank the National *Xenopus* Resource (RRID:SCR013731) and Xenbase (RRID:SCR-003280) for their support. We thank Bret Judson and the Boston College Imaging Core for infrastructure and support. This work was supported by NIH/NIMH (MH109651) to LAL, National Science Foundation IOS award 1656510 to BB, National Institutes of Health grant 8P20GM103449 (Vermont Genetics Network/ Vermont INBRE program) to BB, and National Science Foundation (Grant No. 1626072).

---

**Supplementary Figure 1.**
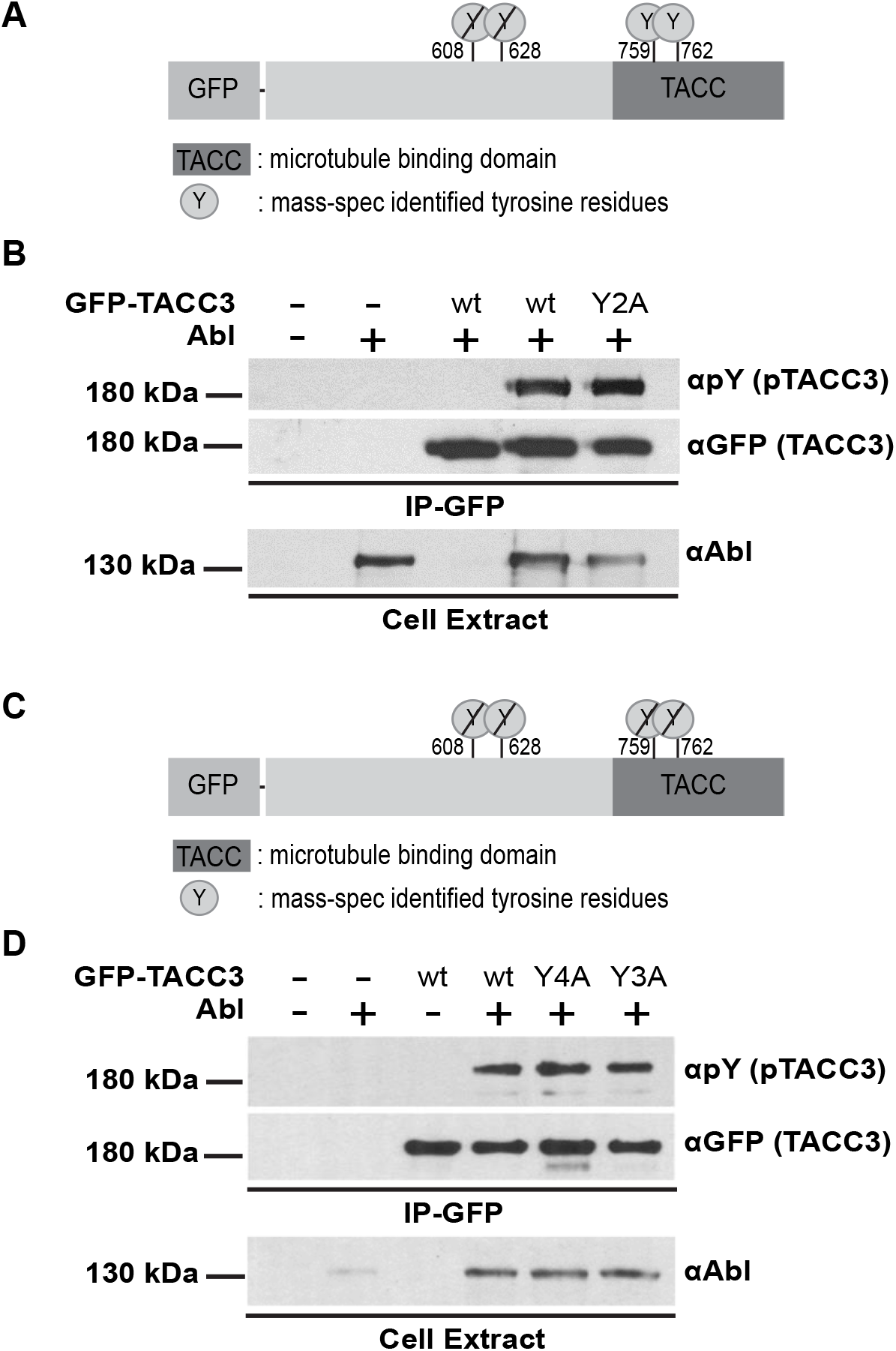
Phospho-null mutations at Mass-spec identified Abelson targeting tyrosine residues do not show reduction in phospho-tyrosine signal levels detected by Western blot. (A,C) Schematic representation of TACC3 full-length protein showing potential tyrosine residues (Y608, Y628, Y759, Y762) targeted for Abelson induced phosphorylation. Combinatorial phospho-null mutants were created for sites Y608 and Y628 (Y2A) or for sites Y608A, Y628A and Y759A (Y3A) or for all tyrosine residues (Y4A), by substituting tyrosine with alanine. (B,D) Western blot performed with phospho-tyrosine specific antibody (pY) showing Abelson induced tyrosine phosphorylation of immunoprecipitated full-length wild-type and phospho-null mutant GFP-TACC3 in the presence of Abelson kinase (top blot). α-GFP shows the expression level of GFP-TACC3 (middle blot) and α-Abl shows the expression of Abelson kinase (bottom blot). Phosphorylation signal is still present to similar extent as compared to wild type (lane 4) when mass-spec identified tyrosine residues are substituted with alanine Y2A, lane 5 (B) Y4A, lane 5; Y3A, lane 6 (D).

**Supplementary Figure 2.**
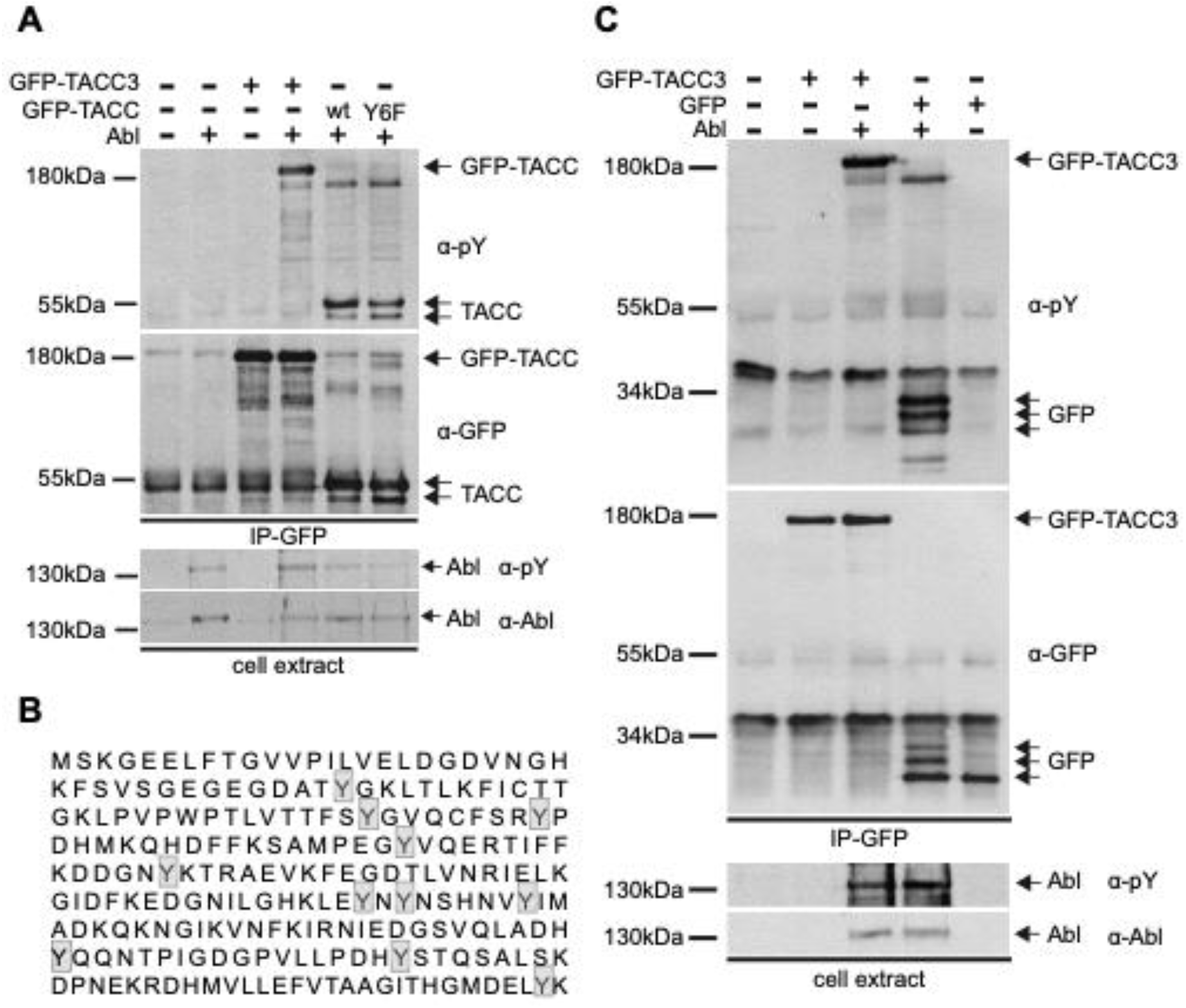
GFP gets phosphorylated in the presence of Abelson kinase and contributes to p-Tyrosine signal obtained with TACC domain tyrosine phospho-null mutant. (A) Western blot performed with phospho-tyrosine specific antibody (αpY, top blot) showing tyrosine phosphorylation levels of immunoprecipitated wild-type full-length GFP-TACC3 with (lane 4, arrow) and without (lane 3, arrow) Abelson kinase and GFP-TACC domain wild-type (lane 5, arrow head) and phospho-null mutant (Y6F, lane6, arrow head) with Abelson kinase. Phosphorylation signal is still present in GFP-TACC phospho null mutant (lane6) to a similar extent as compared to wild type TACC domain (lane 5). 2^nd^ blot from top targeted with α-GFP shows the expression level of GFP-TACC3 and GFP-TACC domain. 3^rd^ blot from top targeted with α-pY shows the phosphorylation levels of Abelson kinase and bottom blot targeted with α-Abl shows the expression of Abelson kinase. IP-αGFP; indicate immunoprecipitation is performed with anti GFP antibody. (B) Amino acid sequence of GFP (UniProtKB/Swiss-Prot: P42212.1), showing tyrosine residues (Y) that are highlighted in a grey box. (C) Western blot performed with phospho-tyrosine specific antibody (αpY, top blot) showing tyrosine phosphorylation levels of immunoprecipitated wild-type full-length GFP-TACC3 with (lane 3) and without (lane 2) Abelson kinase and GFP with (lane 4) and without (lane 5) Abelson kinase. Bands on lane 4 pointed with arrows demonstrating phosphorylated GFP in the presence of Abelson. 2^nd^ blot from top targeted with α-GFP shows the expression of GFP-TACC3 and GFP. 3^rd^ blot from top targeted with α-pY shows the phosphorylation levels of Abelson kinase and bottom blot targeted with α-Abl shows the expression of Abelson kinase. IP-αGFP; indicates immunoprecipitation is performed with anti GFP antibody.

**Supplementary Figure 3.**
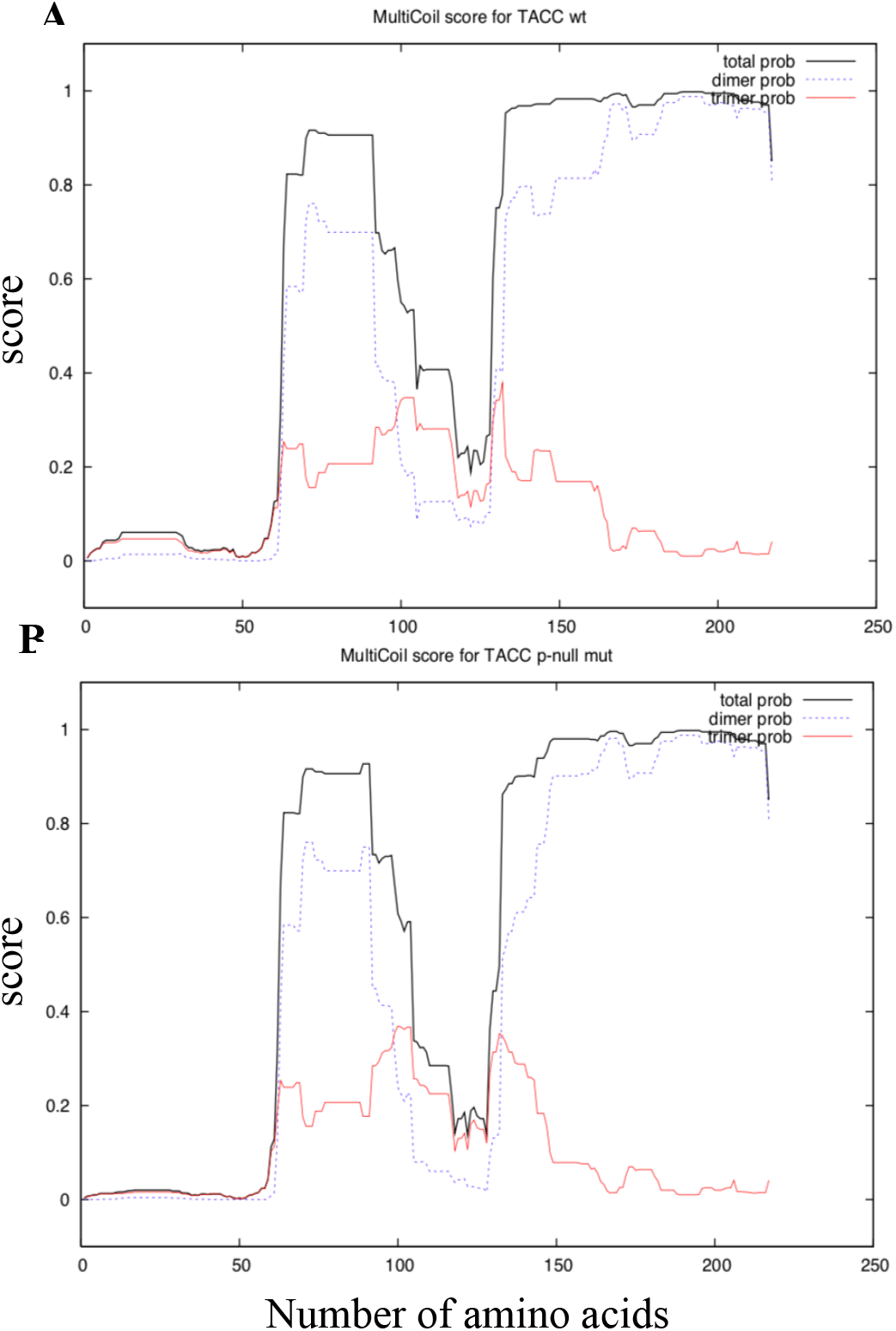
*In silico* prediction of oligomerization of TACC domain wild-type andTACC p-null mutant. (A-B) TACC wt and p-null mutant aminoacid sequence were subjected to coiled-coil and oligomerization prediction using MultiCoil software. Oligomerization probability (score is given on the y-axis) of TACC wild-type is shown at the top panel (A) and TACC p-null mutant is shown at the bottom panel (B). Red line; probability of timerization, dashed line; probability of dimerization, black line; total probability.

**Supplementary Figure 4.**
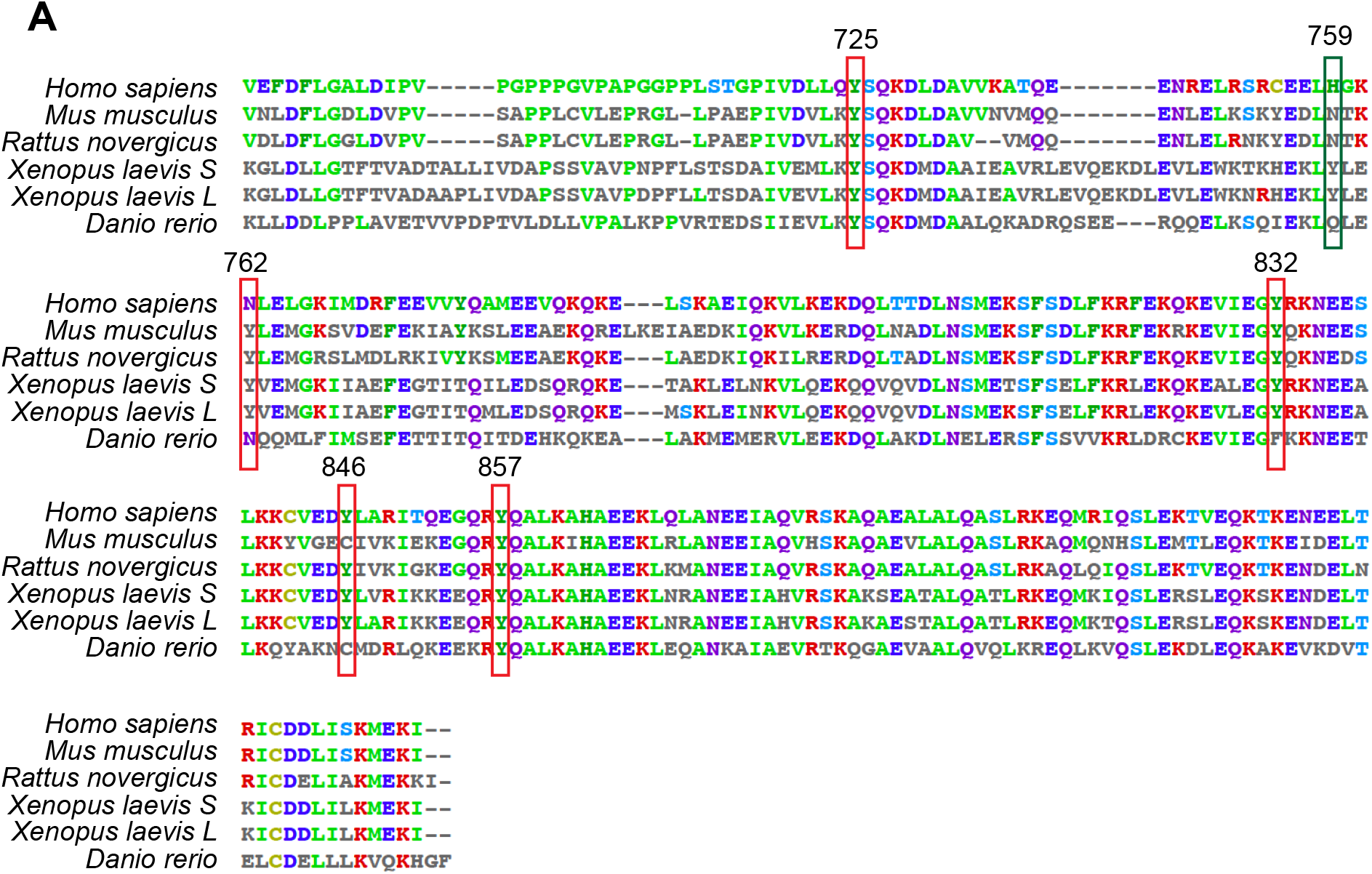
Amino acid sequence alignment of TACC3 showing conservation of tyrosine residues within the TACC domain across species. Tyrosine residues conserved among X. laevis and other species shown in red box. Tyrosine residues that are only present in X. laevis homeologs shown in green box. Accession numbers of the TACC3 sequences aligned; *Homo sapiens* (NP_006333.1), *Mus musculus* (NP_001035525.1), *Rattus novergicus* (NP_001004424.1), *Xenopus laevis* S-homeolog (NP_001081964.1), *Xenopus laevis* L-homeolog (NP_001090422.1), *Danio rerio* (NP_997746.1).

## Supplemental Movie Figure Legends

### Supplemental Movie 1-6

Time lapse live images of cultured *Xenopus laevis* growth cones of GFP-TACC3 wild-type (Movie 1), GFP-TACC3 phospho-null mutants with tyrosine to alanine mutation of tyrosine residues; 608 (Movie 2), 628 (Movie 3), 759 (Movie 4), 762 (Movie 5) and combination of all four tyrosine residues (Y4A, Movie 6). Growth cones are expressing mKate2-MACF43 (magenta) and GFP-TACC3 (green). Images were acquired for 1min with 2 sec intervals. Scale bar, 2μm.

### Supplemental Movie 7-8

Time lapse live images of cultured *Xenopus laevis* growth cones of GFP-TACC domain only wild-type (Movie 7) and GFP-TACC domain phospho-null mutant with all tyrosine residues are mutated into phenylalanine (6xYF, Movie 8). Growth cones are expressing mKate2-tubulin (magenta) and GFP-TACC3 (green). Images were acquired for 1min with 2 sec intervals. Scale bar, 2μm.

### Supplemental Movie 9

Representative phase contrast time-lapse image of a neural tube explant recorded for 4 h with 5 min intervals. Scale bar, 10μm.

### Supplemental Movie 10

Representative phase contrast time-lapse image of a single axon showing the growth track generated by Fiji, Manual Tracking plugin. Tracking data is used to calculate the growth velocity and directionality shown in Figures 4B and F. Image is acquired for 4 h with 5 min intervals. Scale bar, 10μm.

### Supplemental Movie 11

Representative phase contrast time-lapse image of a single axon showing axon pause and retraction. Time-lapse montage of this video is shown in Figure 4C and quantification of the frequency of axon pause and retraction is shown in Figure 4D. Image is acquired for 4 h with 5 min intervals. Scale bar, 10μm.

